# Mathematical model of nuclear speckle morphology

**DOI:** 10.1101/2023.01.12.523856

**Authors:** Shingo Wakao, Noriko Saitoh, Akinori Awazu

## Abstract

Nuclear speckles are nuclear bodies consisting of populations of small and irregularly shaped droplet-like molecular condensates that contain various splicing factors. Recent experiments have shown the following morphological features of nuclear speckles: (I) Each molecular condensate contains SON and SRRM2 proteins, and *MALAT*1 non-coding RNA surrounds these condensates; (II) In the normal interphase of the cell cycle, these condensates are broadly distributed throughout the nucleus in multicellular organisms. In contrast, the fusion of condensates leads to the formation of strongly condensed spherical droplets when cell transcription is suppressed; (III) SON is dispersed spatially in *MALAT1* knocked-down cells, whereas *MALAT1* is dispersed in SON knocked-down cells by the collapse of nuclear speckles. However, the detailed interactions among molecules that reveal the mechanisms of this rich variety of morphologies remain unknown. In this study, a coarse-grained molecular dynamics model of the nuclear speckle was developed considering the dynamics of SON, SRRM2 or SRSF2, *MALAT1*, and pre-mRNA as representative components of condensates. The simulations reproduced the abovementioned morphological changes, by which the interaction strength among the representative components of the condensates was predicted.

## Introduction

Multicellular organisms contain various non-membranous nuclear bodies in the cell nucleus. Their dynamics and biophysical properties are intimately linked to physiological activities, such as gene regulation [1,2]. Typical nuclear bodies, such as the nucleolus and PML bodies, form a spherical shape during interphase, but they change their morphologies under certain cellular or experimental conditions and in diseased cells [1,3].

Nuclear speckles are one of the major nuclear bodies formed by over a hundred protein species, including various pre-mRNA splicing factors [4,5,6] and noncoding RNAs, like *MALAT1* [7,8]. Nuclear speckles are populations of small irregular-shaped droplet-like molecular condensates distributed throughout the nucleus, enriched in the inter-chromosome compartments [5,9,10,11,12,13,14,15,16]. Transcriptionally active genes are often found at the nuclear speckle periphery [11], and their transcripts are associated with or transit through nuclear speckles [17,18].

Nuclear speckles were originally identified via immunofluorescence using the monoclonal antibody mAb SC35 that was raised against biochemically isolated mammalian spliceosomes [19]. This antibody had been long considered to be based on immunoblots, and that it recognizes a pre-mRNA splicing factor of 35 kDa of SRSF2/SC35, with possible cross-reactivity to other splicing factors harboring repetitive serine and arginine (SR) sequences. A recent study re-characterized mAb SC35, revealing that its main target may be SRRM2 (formally known as SRm300), a spliceosome-associated protein [15]. Because the antigens remain somewhat elusive, we herein refer proteins probed with mAb SC35 as SC35p, which stands for mAb SC35-recognizing protein candidates. Among these, we have focused on SRRM2 and SRSF2. Recent studies involving high-resolution fluorescence imaging reported the following morphological features of nuclear speckles: I) During interphase, the core region of each small molecular condensate of nuclear speckles contained SON and SC35p, and *MALAT1* tended to surround them like a shell (Figure 1A) [20]; II) When all gene transcriptions were suppressed, the molecular condensates fused with each other and formed dense spherical condensates with stronger fluorescent intensity than those in normal interphase (Figure 1B) [13,21,22]; III) Conversely, when *MALAT1* was knocked-down, SC35p was sustained in the condensates but SON dispersed (Figure 1C) [20]; IV) Similarly, when SON was knocked-down, SC35p was sustained in the condensates but *MALAT1* dispersed (Figure 1D) [8,20,21]. In the latter two cases, nuclear speckles were collapsed.

**Figure 1.**
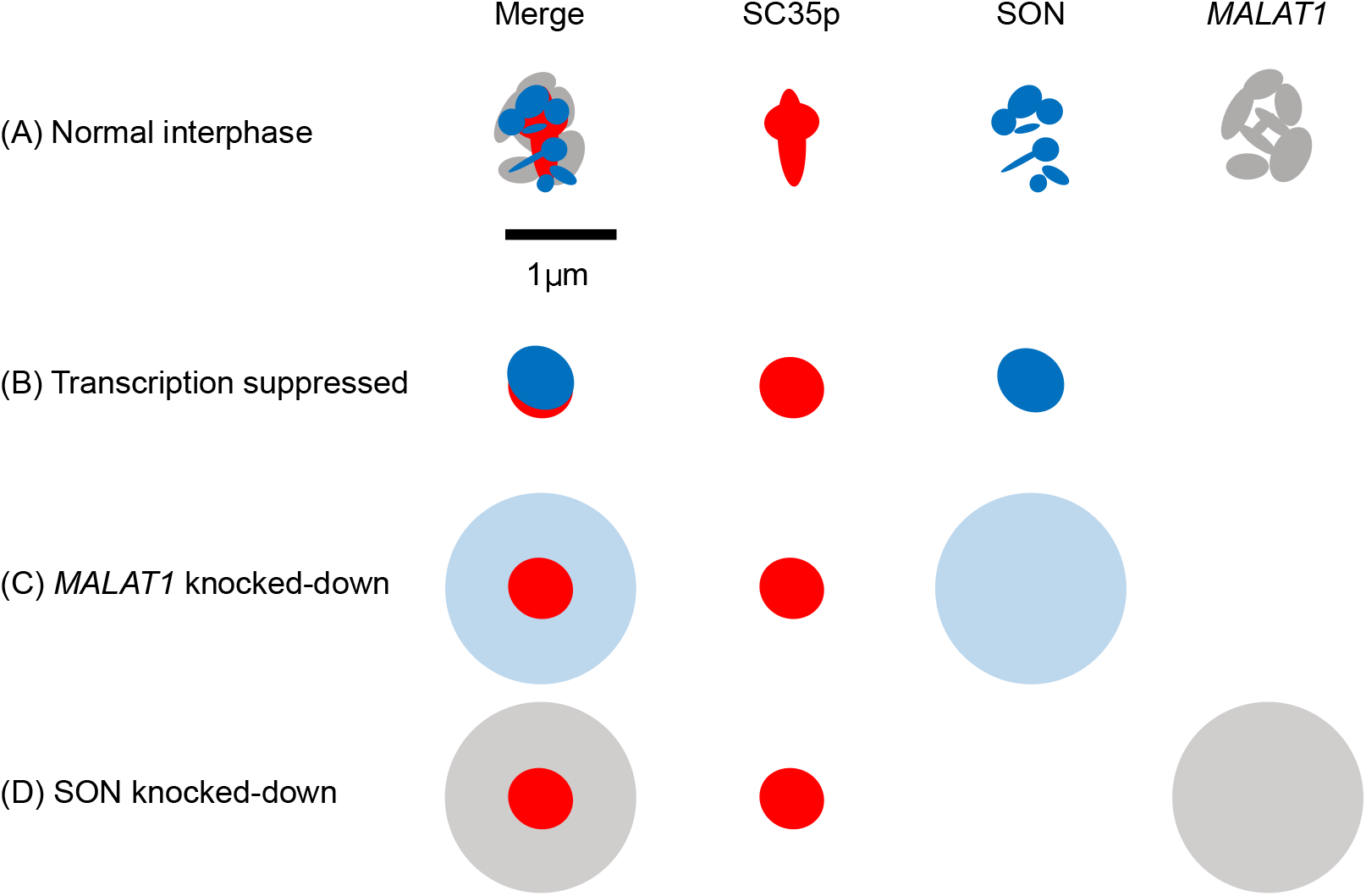
Illustrations of molecular distributions in the local region of nuclear speckles observed using fluorescence microscopy imaging under various conditions. Distributions of SC35p, SON, and *MALAT1* in (A) normal interphase, (B) transcriptional repression state, (C) *MALAT1* knocked-down state, and (D) SON knocked-down state [8,13,20,21,22].

The representative molecules regulating nuclear speckle morphology were identified via the aforementioned experiments. However, the detailed interactions among these molecules that could reveal the underlying mechanisms remain unclear. For other nuclear bodies, such as the nucleolus and paraspeckle, the development of mathematical models provided deeper insights into the relationships between their morphologies and molecular chemo-mechanical features [3,23,24]. Recently, a mathematical model of the molecular condensates of nuclear speckles was developed that accounted for the interactions among SON, SC35p, and *MALAT1* and reproduced their hierarchical condensations [20]. However, this model had some flaws. For example, it was easily expected that the dispersion of SON by *MALAT1* knockdown could not be reproduced because this model assumed the SON-SON attractive forces as well as SON-SC35p attractive forces.

In this study, we aimed to reveal the mechanism and role played by each molecule of the nuclear speckle in determining its morphological features. To that end, we developed a coarse-grained molecular dynamics model of the local region of nuclear speckles with a few molecular condensates. SON, SC35p, *MALAT1*, and pre-mRNA were considered as the representative components of droplets. Simulations of the proposed model with appropriate assumptions of the effective interactions among these molecules reproduced all the above-mentioned morphological changes in the local region of the nuclear speckle. The results also suggested that SRRM2 and *MALAT1* were regarded as the core and shell components of core-shell structures in nuclear speckles.

## Materials and Methods

### Candidates of Proteins Probed with mAb SC35 (SC35p): SRSF2 and SRRM2

mAb SC35 has been used to observe nuclear speckles as this antibody recognizes SRSF2/SC35. However, a recent re-characterization study revealed that a major target for this antibody was actually SRRM2 [15]. Therefore, in this study, SRSF2 and SRRM2 were assumed as SC35p for their individual models. The model for assuming SC35 as SRSF2 was called the “SF2 model”, and that for assuming SC35 as SRRM2 was called the “RM2-model”.

### Coarse-Grained Models of SON, SRRM2, and SRSF2

All three focused proteins, SON, SRRM2, and SRSF2, are intrinsically disordered proteins that involve partially folded regions, and their intrinsically disordered regions (IDRs) occupy a large ratio of their amino acid sequences (Figure 2A, B) [4,25,26]. This indicates that they do not possess unique stable folding structures but change their shapes like random polymers. Therefore, each of these proteins was described using one soft spherical particle with their respective radii *r*^*p*^ (*p*: SON,SRRM2,SRSF2) for simplicity. Here, the particles representing SON, SRRM2, and SRSF2 were named SON, SRSF2, and SRRM2 particle, respectively. Each particle indicates the spatial region that the chain on the protein tends to occupy. Therefore, the excluded volume effects among these particles were assumed to be so weak that a small ratio of the regions in the two contacting particles was allowed to overlap with each other.

**Figure 2.**
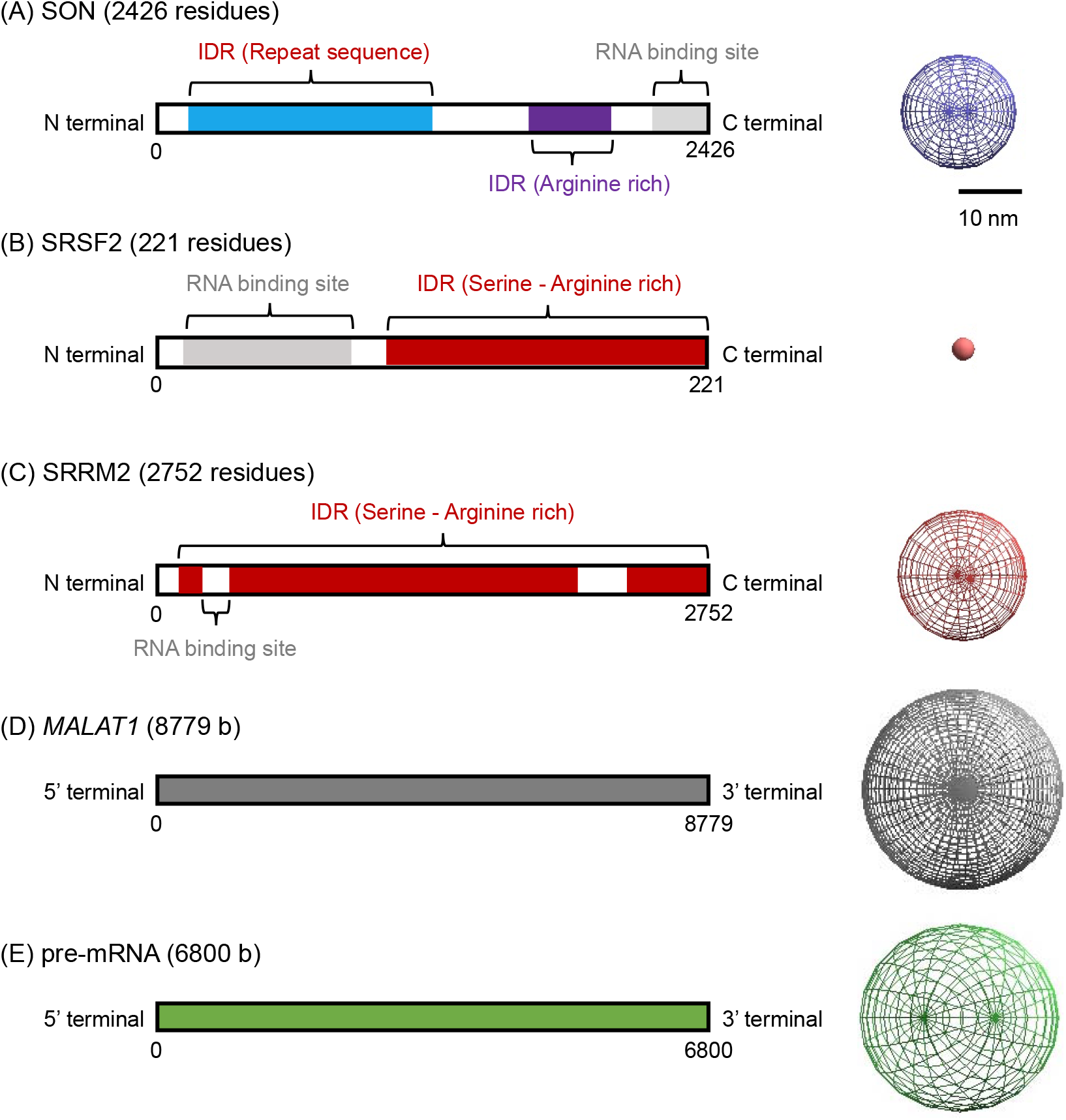
Construction of a coarse-grained model of each molecule. Lengths and local regions in proteins and RNAs (left) and soft spherical particle models (right) of (A) SON, (B) SRSF2, (C) SRRM2, (D) *MALAT*1, and(E) pre-mRNA. IDR, intrinsically disordered region.

The radii *r*^*p*^ were assumed according to the arguments of random polymers with excluded volume as 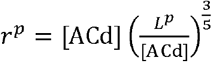, where [ACd] = 0.36 (*nm*) indicates the average distance between the edge of the amino group and that of the carboxyl group in each amino acid, and *L*^*P*^ indicates the effective length of proteins. In the present study, for the SON and SRSF2 models, *L*^*p*^ was defined as the edge-to-edge distance of proteins containing secondary structures along the, amino acid sequence as follows: *L*_*P*_ = [sum of all distances between N-terminals and C-terminals of partially folded regions (*nm*)] + [ACd] · [Number of amino acids belonging to IDR] (Figure 2A, B, Supplementary Figure S1). The three-dimensional coordinates of atoms and positions of folded regions in SON and SRSF2 were obtained in PDB format files from the UniProt database [https://www.uniprot.org] ID Q01130 and P18583 [27], where these features were inferred via Alpha-Fold2 [28,29]. Based on these estimates, *r*^SON^ = ∼ 18.4 nm and *r*^SRSF2^ = ∼ 3.4 nm were obtained.

In this study, SRRM2 was assumed to be a simple unfolded random polymer of amino acids because most of the regions were IDR, and no public data of the three-dimensional coordinates of atoms and positions of folded regions in SRRM2 were available (Figure 2C). Therefore, LP = [ACd] · [Number of amino acids belonging to IDR] was assumed, which gave *r*^SRRM2^ = ∼ 20.8 nm.

In the present cases, the rigidity of the random polymer is known to decrease with an increase in the number of segments corresponding to 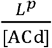. The focused molecules were expected to be more rigid than random polymers in their central regions but more similar to them in their outer regions.

Therefore, in this model, the rigidity parameters of these particles, *q*^*p*^, were assumed to be proportional to the estimated conventional arguments of the rubber elasticity of ideal chains, as follows: 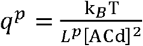. Recent molecular dynamics simulations showed that the elastic stress-strain relations of random polymers with excluded volumes could be approximated by those of ideal chains when the polymer concentrations were not as high as those of solids [30]. Here, *q*^SON^ = ∼ 1.43×10^−5^ kg · s^−2^, *q*^SRRM2^ = ∼ 1.16×10^−5^ kg· s^−2^, and *q*^SRSF2^ = ∼ 2.35×10^−4^ kg · s^−2^ were obtained.

### Coarse-Grained Models of *MALAT1* and Pre-mRNAs

In the present model, RNAs were assumed to be unfolded random polymers, and each RNA chain was described as one soft spherical particle with their respective radii *r*^R^ (*R*: *MALAT1* or pre-mRNA). Here, the particles representing *MALAT1* and pre-mRNA were respectively called *MALAT1* and pre-mRNA particles. Similar to protein models, each particle indicated the spatial region that the chain on the protein tended to occupy (Figure 2D, E); therefore, the excluded volume effects among these particles were assumed to be so weak that a small ratio of the regions in the two contacting particles were allowed to overlap with each other.

The radii *r*^*R*^ were assumed as 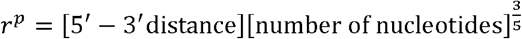 where [5^′^ 3^′^distance] was assumed as 0.3 nm, and the number of nucleotides for *MALAT1* was assumed as 8779 bp. In the present study, the number of pre-mRNAs was assumed to be a unique value of 6.8 mb that was the same as the average base pairs of gene-transcribed regions in the human genome, estimated based on the Refseq data [31]. The rigidity parameters of these articles, *q*^*R*^, were also estimated in the same way as the above-mentioned cases of proteins, where 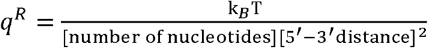 was assumed. Further, *r*^MALAT1^ = ∼ 33.0 nm, *r*^pre-mRNA^ = ∼ 29.9 nm, *q*^MALAT1^ = ∼ 5.75× 10^−6^ kg · s^−2^, and *q*^pre-mRNA^ = ∼ 6.76 · 10^−6^ kg · s^−2^ were obtained.

### Motion of Each Molecule

Because each biomolecule was expected to exhibit three-dimensional Brownian motion in the nucleus, the position of the *i*-th particle with radius, in the (*x,v,z*) three-dimensional space, *x*_*i*_, = (*x*_*i*_, *y*_*i*_,*z*_*i*_), was assumed to obey the Langevin equation as follows:

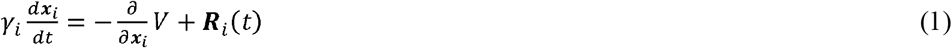

where *γ*_*i*_ and **R**_*i*_(*t*) are the coefficients of drag force and Gaussian white noise, respectively, playing the role of the random force from the nucleoplasm to *i*-th particle, obeying ⟨**R**_*i*_(*t*) ⟩ = 0 and ⟨**R**_*i*_(*t*) **R**_*i′*_ (*t*) ⟩) = 6*γ*_*I*_ *k*_*B*_ *Tδ*_*i i′*_*δ*(*t* − *s*), with Boltzmann constant *k*_*B*_ and temperature *T*, where *k*_B_*T* = 4.141947×10^−21^ kg m^2^ s^-2^ (*T* = 300 K). Here, *δ*_*ij*_ indicates the Kronecker delta and *δ*(…) indicates the Dirac delta function. The term *V* indicates that the potential of the system involves the potential of forces among the particles and effect of the boundary conditions. The drag coefficient y for each particle was assumed as 6*πηmr*_*i*_, with the viscosity of the nucleoplasm *η* = 0.64 *kg m*^−1^ s^−1^ [32].

Particle motion was assumed to be restricted to a spherical region with a radius of 500 *nm*. Note that 500 nm was slightly larger than the characteristic length of the inter-chromosome components in the cell nucleus. Such a wide space was employed in the simulations to reveal the mechanism of the droplet morphologies from the self-assembling behaviors of nuclear speckle components without the influence of other factors, such as chromatin territories and other nuclear bodies.

### Model of Effective Interactions among Molecules

The potential of system *V* to provide the forces working on and among the particles is given as:

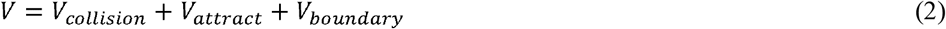

where *V*_*col*_ is the potential of the collisional interactions among the particles owing to their excluded volumes, *V*_*attract*_ is the potential of the attractive interactions among the particles owing to the electrostatic interactions or biochemically specific affinities among molecules, and *V*_*boundary*_ is the potential of the effect of the boundary wall to restrict particles in the above-mentioned spherical region.

The potential *V*_*col*_ was assumed via the following:

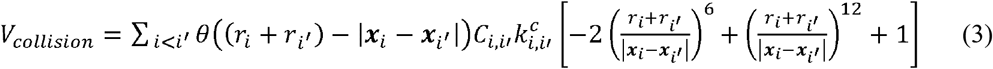

where 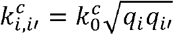 represents the repulsion strength, *C*_*i,i*′_ is the value of *C*_*i,i*′_ = 1 (0) when repulsion works (does not work) in between *i*-th and *i*-th particles.

*θ* is the Heaviside step function, defined as follows:

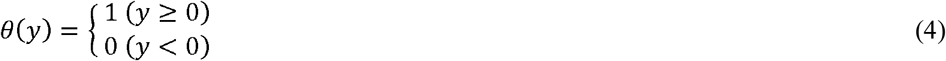

The potential *V*_*attract*_ was assumed by the following equation:

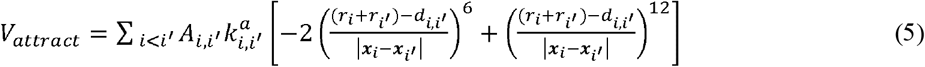

where 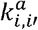 represents the strength of the attractive force, and *A*_*i,i′*_ is the value that *A*_*i,i′*_ =1 (0) when attractive force works (does not work) in between *i*-th and *i*-th particles. *d*_*i,i′*_ was assumed to be smaller than *r*_*i*_ + *r*_*i*′_ because the particles representing the space occupied by polymers were assumed to be so soft that the stable distance between two of them was expected to be nearer than the sum of their radii when attractive forces work in between them. In this study, *d*_*i,i′*_ was chosen to satisfy the following relation:

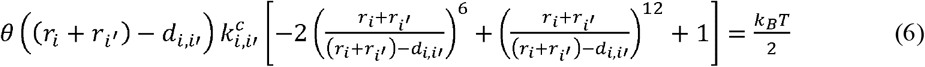

where *d*_*i,i′*_ provided the typical length of the region where two particles could overlap when only the repulsion represented by Eq. (3) was assumed to work in between them (Supplementary Figure S2).

The potential *V*_*boundary*_ was assumed by the following equation:

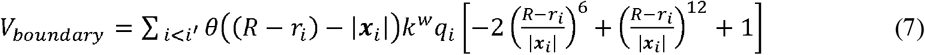

where *k*^*w*^ represents the strength of the restriction from the boundary wall, and *R* represents the radius of the simulation space, assumed as 500 *nm*.

### Simulation and Data Analysis Method

The time integral of Langevin Eq. (1) was calculated numerically using the Eular-Maruyama method with a unit step of 3.16 × 10^−7^ (sec). For the initial condition (time = 0 (*sec*)), the particles were placed randomly in the sphere within a radius of 250 *nm*. The radial distributions of distances between two particles are defined as *R*^*X,Y*^(*dist*) = [Probability that the distance between X and Y particles = *dist*]. Note that the accumulation or dispersion features of the X and Y particle populations were provided by the profile, in particular the height, width, and tail of the first peak nearest from *dist* = 0 of *R*^*X,Y*^(*dist*).

Recent experimental studies suggested that SC53p always condensates independently of the cell state [8,13,20,21,22]. Therefore, to evaluate the morphological features of particle condensates, the radial distributions of SC53p (SRRM2 or SRSF2), SON, *MALAT1*, and pre-mRNA particles from SC53p (SRRM2 or SRSF2) particles were measured; the radial distribution of X particle (X = SON, SRSF2, SRRM2, *MALAT1*, or pre-mRNA) was defined as *R*^*X*^(*dist*)= [Probability that the distance between X and SC53p (SRRM2 or SRSF2) particles = d,sL]. Additionally, the from other *MALAT1* particles is defined as *R*^*SS*^ (*dist*) = [Probability that the distance between a radial distributions of SON particles from other SON particles and those of *MALAT1* particles SON and other SON particles = *dist*] and *R*^*MM*^(*dist*) = [Probability that the distance between a *MALAT1* and other *MALAT1* particles = *dist*], which were also measured to evaluate their between the time interval 403– 474 (*sec*), where the radial distribution of the typical state was condensations or dispersions. These distributions were estimated using the particle configurations obtained without the influence of the initial particle configuration.

## Results

### Simulation of RM2 Model using Appropriately Designed Parameters of Specific Molecular Interactions Reproducing Droplet Morphologies in Nuclear Speckles

The RM2 model with the following specific intermolecular interactions was simulated (Figure 3A, Supplementary Figure S2A), which could reproduce the experimentally observed morphological features of droplets of nuclear speckles (Figure 1):

**Figure 3.**
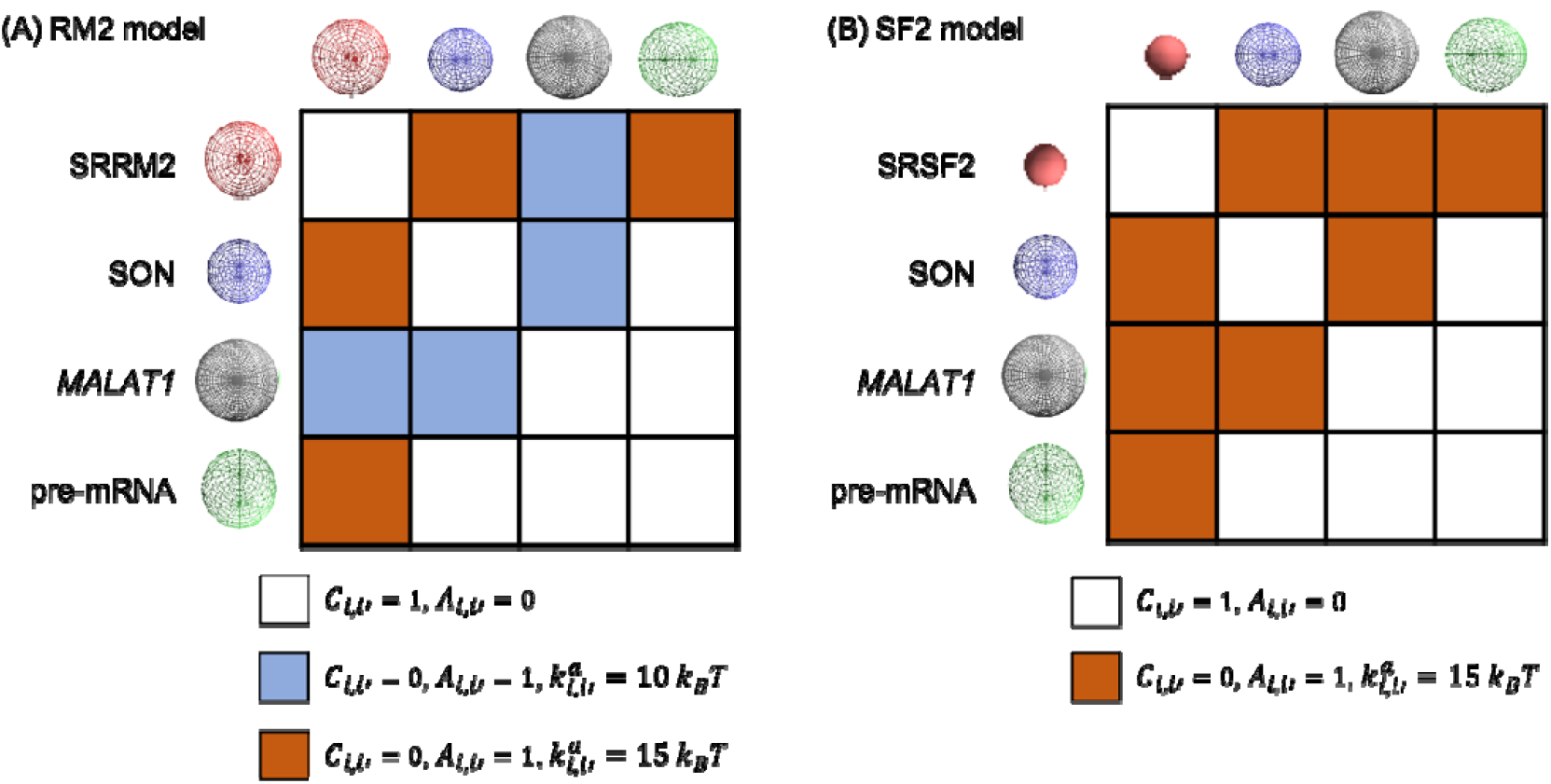
Matrices of interaction networks among particles in (A) RM2 and (B) SF2 models with appropriately designed parameters. White boxes indicate repulsion, blue boxes indicate weak attraction, and brown boxes indicate strong attraction between a pair of particles. Each profile of interaction potential between each pair of particles is shown in Supplementary Figure S2.

I. Repulsive forces caused by the excluded volumes of molecules between two particles representing the same molecular species were assumed as *C*_*i,i′*_ =1 and *A*_*i,i′*_ = 0. The parameter providing the strength of the repulsive forces, 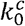 was given as 1.4×10^3^ *k*_*B*_*T*.
II. A specific attractive force between SRRM2 and SON particles and between SRRM2 and pre-mRNA particles was assumed as *C*_*i,i′*_ = 0, *A*_*i,i′*_ = 1. The attractive strength II) parameter was assumed uniformly as 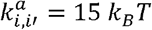.
III. The specific attractive force between SRRM2 and *MALAT1* particles and between SON and *MALAT1* particles was assumed as *C*_*i,i′*_ = 0, *A*_*i,i′*_ = 1. The attractive III) strength parameter was assumed uniformly as 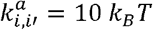.
IV. The interaction between SON and pre-mRNA particles and that between *MALAT1* and pre-mRNA particles were assumed to be repulsive, as *C*_*i,i′*_ = 1, *A*_*i,i′*_ = 0, and 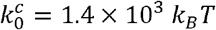.

First, the numbers of SRRM2, SON, *MALAT1*, and pre-mRNA particles were 60, 60, 60, and 420, respectively, and was simulated to consider the behavior of droplets of nuclear speckles under normal conditions during interphase. The present model exhibited broadly distributed particle condensates (Figure 4A). Further, *R*^*SON*^(*dist*), *R*^*SRRM2*^(*dist*), and *R*^*MALAT1*^(*dist*) exhibited a peak at the similar near, but involved a lower peak and longer tail than and. This fact indicated that the condensates of SRSF2 and SON particles were surrounded by *MALAT1* particles similar to the core-shell structure observed in experiments (Figure 1A) [14,20].

**Figure 4.**
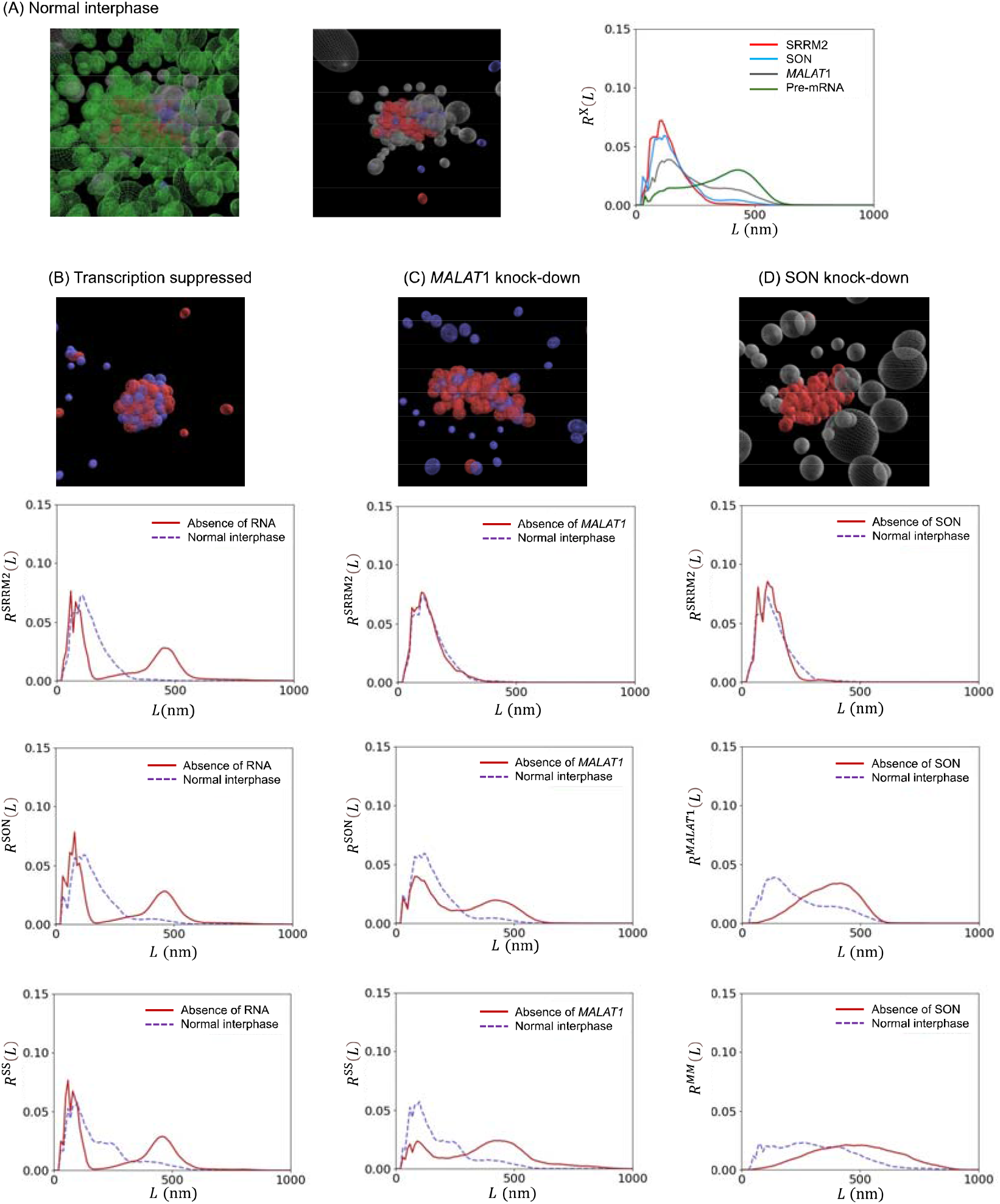
Simulation results of the RM2 model with appropriately designed parameters. (A) Typical snapshots of the simulations (left), the same snapshots but without pre-mRNA (middle), and radial distributions of particles, *R*^SRRM2^(*L*), *R*^SON^(*L*), *R*^*MALAT1*^(*L*), and *R*^pre-mRNA^(*L*) (right) in the case of normal interphase. (B–D) Typical snapshots of the simulations without pre-mRNA (first), *R*^SRRM2^(*L*) (second), *R*^SON^ (*L*) or *R*^*MALAT1*^(*L*) (third), and *R*^SON^(*L*) or *R*^*MALAT1*^ (*L*) and *R*^SS^(*L*) or *R*^*MM*^(*L*) (fourth), in the case that (B) transcription was suppressed, (C) *MALAT1* was knocked-down, and (D) SON was knocked-down. Each particle color indicates molecular species as explained in Figure 2.

Second, to consider the situation after transcription activity was suppressed in the entire nucleus, the numbers of SRRM2, SON, *MALAT1*, and pre-mRNA particles were 60, 60, 0, and 0, respectively. The positions of the large peaks that were the nearest from of, and had shifted to close to, and their tails became shorter than the previous case (Figure 4B). This indicates that SRRM2 and SON had formed a denser condensate than the previous case, in which denser droplets were formed owing to the transcription suppression observed in the experiments (Figure 1B) [13,21,22].

Third, to consider the *MALAT1* knocked-down state, the numbers of SRRM2, SON, *MALAT1*, and pre-mRNA particles were 60, 60, 0, and 420, respectively. In this case, the qualitative features of were almost unchanged from those in the first case. However, and were broader, and the peak nearest to was much smaller than that in the first case (Figure 4C). This indicates that SON was spatially dispersed, as observed in *MALAT1* knockdown experiments (Figure 1C) [20].

Finally, to consider the SON knocked-down state, the numbers of SRRM2, SON, *MALAT1*, and pre-mRNA particles were 60, 0, 60, and 420, respectively. In this case, the qualitative features of were almost unchanged from those in the first case. However, and were broader, and the peak nearest to was much smaller than that in the first case (Figure 4D). This indicates that *MALAT1* was spatially dispersed, as observed in the SON knockdown experiments (Figure 1D) [8,20,21].

### Simulation of SF2 Model using Appropriately Designed Parameters of Specific Molecular Interactions Reproducing Droplet Morphologies in Nuclear Speckles

The SF2 model could also exhibit the following morphological features of particle condensates similar to the experimentally observed droplets of nuclear speckles (Figure 1) when the appropriate specific intermolecular interactions were simulated (Figure 3B, Supplementary Figure S2B):

I. The interactions among SON, SRSF2, and pre-mRNA were assumed to be the same as those among SON, SRRM2, and pre-mRNA particles assumed in the previous RM2 model. The interaction between *MALAT1* and pre-mRNA particles was assumed to be the same as in the previous SF2 model.
II. Specific attractive forces were assumed between SRSF2 and *MALAT1* particles and between SON and *MALAT1* particles as C_*i,i′*_ = 0, *A*_*i,i′*_ = 1, and 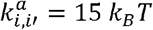.

Note that the appropriate interaction strengths between *MALAT1* particles and other particles seemed to change slightly, which may have been owed to different particle sizes between SRSF2 and SRRM2 particles. Additionally, the number of SRSF2 particles was assumed as twice that of SON and *MALAT1* particles as referred to the previously proposed mathematical model [20], where the number of SRSF2, SON, MALAT1, and pre-mRNA particles were assumed as 120, 60, 60, and 420, respectively. With these assumptions, the SF2 model could qualitatively exhibit similar behaviors of SON, SC35p, and *MALAT1* particles to those obtained in the previous RM2 model except for the following facts (Figure 5, Supplementary Figure S3).

**Figure 5.**
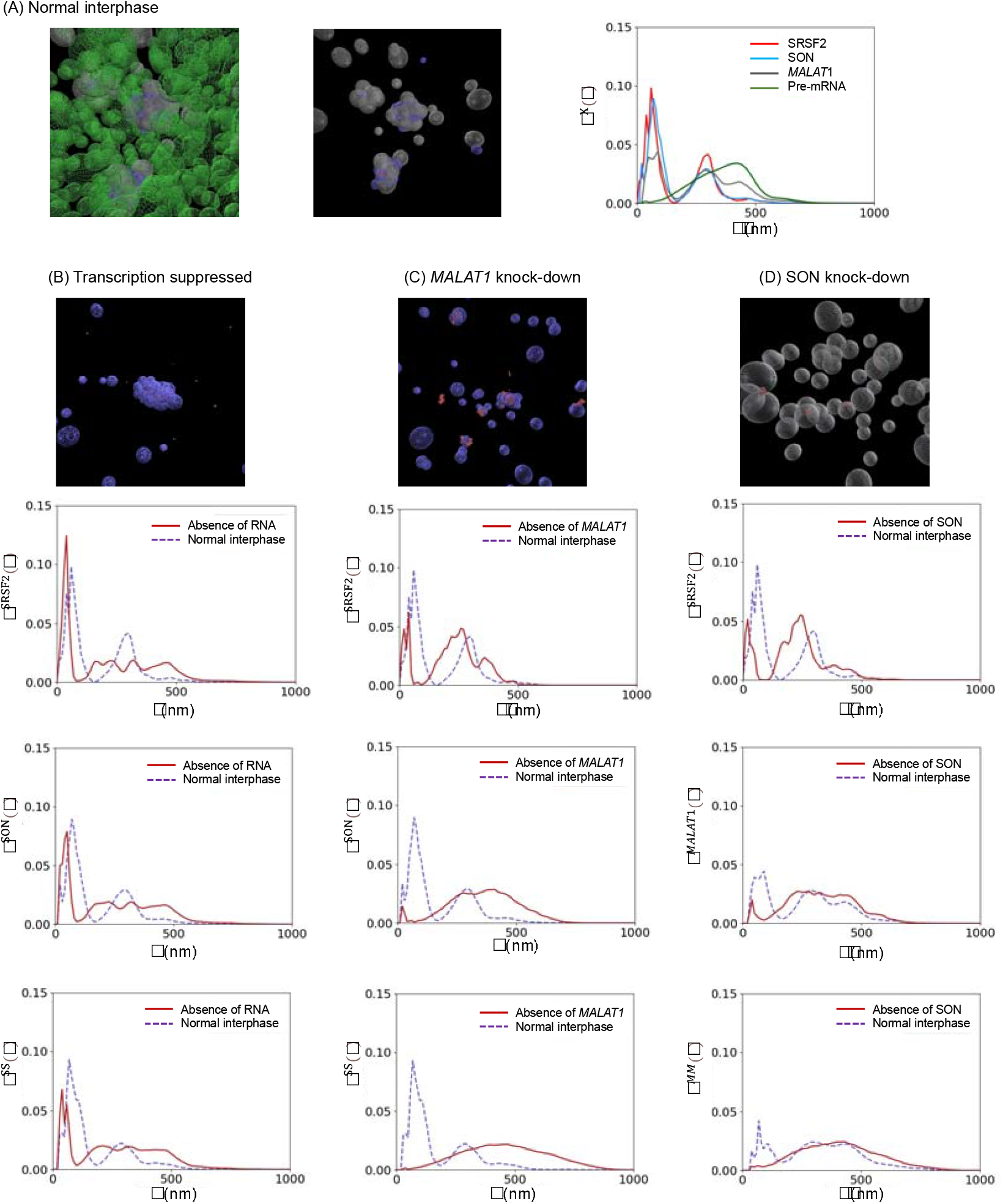
Simulation results of the SF2 model with appropriately designed parameters. (A) Typical snapshots of the simulations (left), the same snapshots but without pre-mRNA (middle), and radial distributions of particles, *R*^SRSF2^(*L*), *R*^SON^(*L*), *R*^*MALAT1*^ (*L*), and *R*^pre-mRNA^(*L*) (right) in the case of normal interphase. (B–D) Typical snapshots of the simulations without pre-mRNA (first), *R*^SRSF2^(*L*) (second), *R*^SON^(*L*) or *R*^*MALAT1*^(*L*) (third), and *R*^SON^(*L*) or *R*^*MALAT1*^ (*L*) and *R*^SS^(*L*) or *R*^*MM*^(*L*) (fourth), in the case that (B) transcription was suppressed, (C) *MALAT1* was knocked-down, and (D) SON was knocked-down. Each particle color indicates molecular species as explained in Figure 2.

i. In the normal interphase simulation, *R*^*SON*^ (*dist*), *R*^*SRSF2*^(*dist*), and *R*^*MATAL1*^(*dist*) exhibited similar profiles near *dist* = 0, and *R*^*MATAL1*^(*dist*) did not show a longer tail (Figure 5A) than in the RM2 model (Figure 4A). Furthermore, SRSF2 and SON particles were not surrounded by *MALAT1* particles, which differed from the core-shell structure observed in experiments [14,20].
ii. In the *MALAT1* knockdown and SON knockdown simulations, the large peaks of *R*^*SRSF2*^ (*dist*) near *dist* = 0 appeared but their heights became lower (Figure 5C–D), which differed from those in the RM2 model (Figure 4C–D). These facts indicated that the SRSF2 condensation was also weakened because of MALAT1 or SON knockdown, which differed from past experiments [8,20,21].

### Simulations using Interaction Parameters Assumed by Electrostatic Properties of Molecules

Regarding the electrostatic properties of molecules, the interaction parameters between each pair of particles can be assumed as follows:

The IDRs of SRRM2 contained 642 serine and 468 arginine residues, and those of SRSF2 contained 42 serine and 35 arginine residues. Therefore, SC35p was expected to exhibit a negative charge. Conversely, SON was expected to exhibit a positive charge because the serine-arginine-rich IDR contains more number of arginine residues (n = 87) than serine residues (n = 67). Therefore, *C*_*i,i′*_ =0 and *A*_*i,i′*_ =1 may be assumed between the SC35p and SON particles because the electrostatic attractive forces may work in between these particles (Figure 6A, B). Conversely, *C*_*i,i′*_ =1 and *A*_*i,i′*_ =0 may be assumed between two SRRM2 particles, two SRSF2 particles, and two SON particles because both particles in each pair have the same charges (Figure 6A, B).

**Figure 6.**
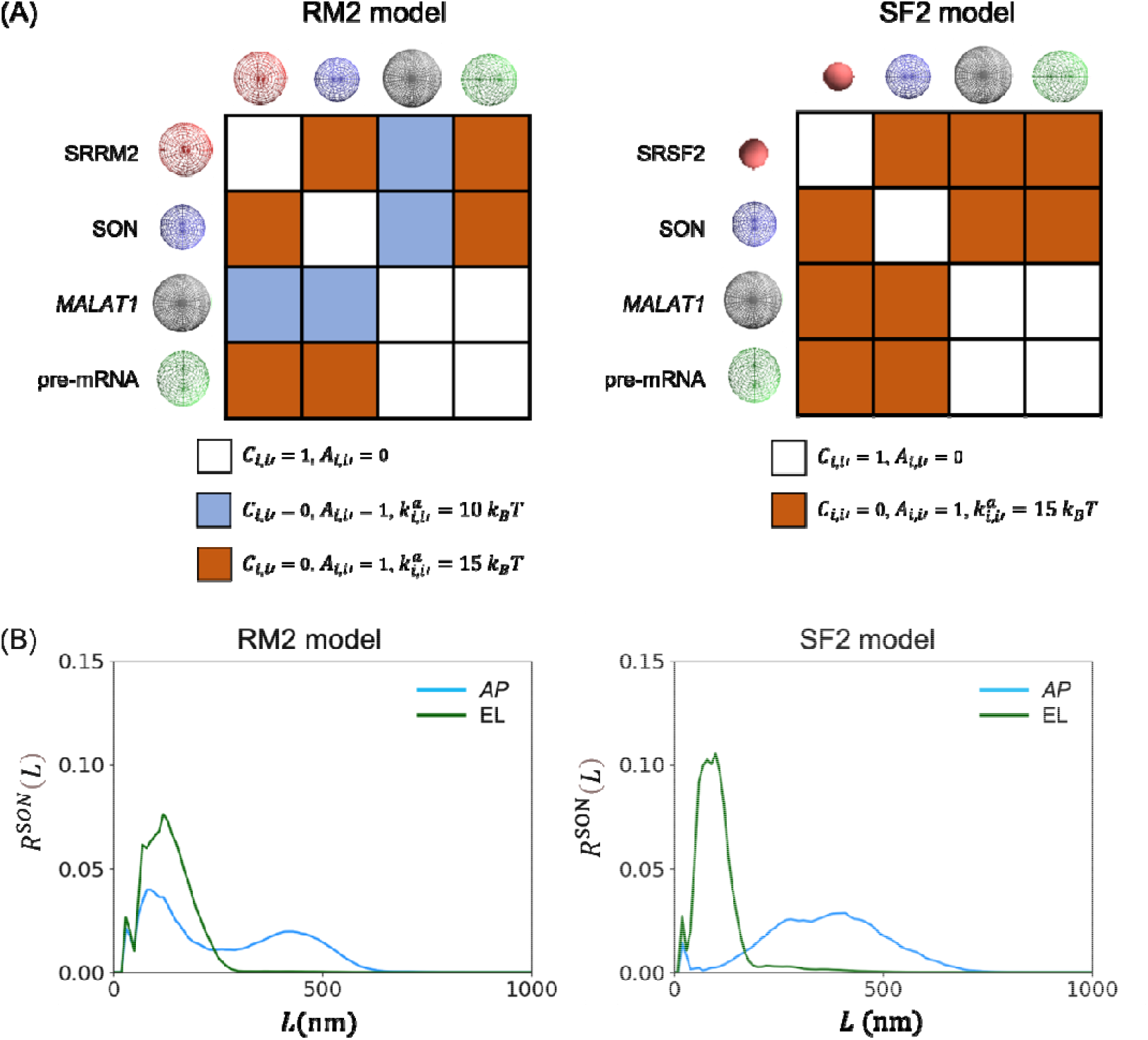
Simulation results of RM2 and SF2 models with parameters based on electrostatic properties. (A) Matrices of interaction networks among particles. White boxes indicate repulsion, blue boxes indicate weak attraction, and brown boxes indicate strong attraction between a pair of particles. (B) Radial distributions of SON particles in RM2 and SF2 models with appropriately designed parameters (AP) and electrostatic property-based parameters (EL) in the *MALAT1* knockdown state.

RNAs are negatively charged polymers. Therefore, *C*_*i,i′*_ =1 and *A*_*i,i′*_ I =0 were assumed between two *MALAT1* particles, two pre-mRNA particles, and *MALAT1* and pre-mRNA particles (Figure 6A).

Both SRRM2 and SRSF2 were expected to be involved in RNA-binding sites [27]. The binding of two molecules through such specific binding sites is generally expected to be stronger than electrostatic forces. Therefore, although the total charges of SRRM2 and SRSF2 were assumed to be negative, *C*_*i,i′*_ and *A*_*i,i′*_ =1 could be assumed between SC35p and RNA (*MALAT1* and pre-mRNA) particles (Figure 6A, B).

Similar to SC35p, SON also contains RNA-binding sites [27,33]. Additionally, the total charge of SON was expected to be positive. Therefore, an attractive force between the SON and RNA particles was naturally assumed as *C*_*i,i′*_ =0 and *A*_*i,i′*_ = 1. Here, the attraction strength between SON and RNA particles was expected to be smaller than that between SON and SC35p, because RNAs were expected to have a stronger negative charge than SC35p.

Notably, the assumptions of the interactions among particles in the previous section were consistent with the above-mentioned intuitive molecular interactions based on electrostatic forces, except for the interactions between SON and pre-mRNA. Therefore, the RM2 and SF2 model simulations were performed using the same interaction parameters among molecules as in the previous arguments, except for those between SON and pre-mRNA, where *C*_*i,i′*_ = 0, *A*_*i,i′*_ = 1, and 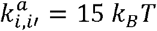 were assumed. In these cases, SON dispersion did not occur even when the model was simulated without *MALAT1* (Figure 6C, D), indicating that the spatial dispersion of SON due to *MALAT1* knockdown (Figure 1C) could not be reproduced in either model.

## Discussion

In this study, a coarse-grained molecular dynamics model, with SON, SC35p (SRRM2 or SRSF2), *MALAT1*, and pre-mRNA as the representative components of condensates, was developed to simulate the morphological behavior of droplet-like condensates of nuclear speckles. Simulations of the model assuming SRRM2 as SC35p reproduced various experimentally observed cell state-dependent droplet morphologies. Simulations of the model assuming SRSF2 as SC35p also exhibited similar behaviors but some features were inconsistent with experimentally observed phenomena. These facts suggested that SC35 should be SRRM2 rather than SRSF2, supporting a recent proposal [15]. Additionally, the effective interactions and their strengths among the representative components of nuclear speckles were predicted through these simulations.

Our findings suggest the following mechanism for the cell state-dependent morphology of nuclear speckles as well as the roles played by the representative molecules (Figure 7):

**Figure 7.**
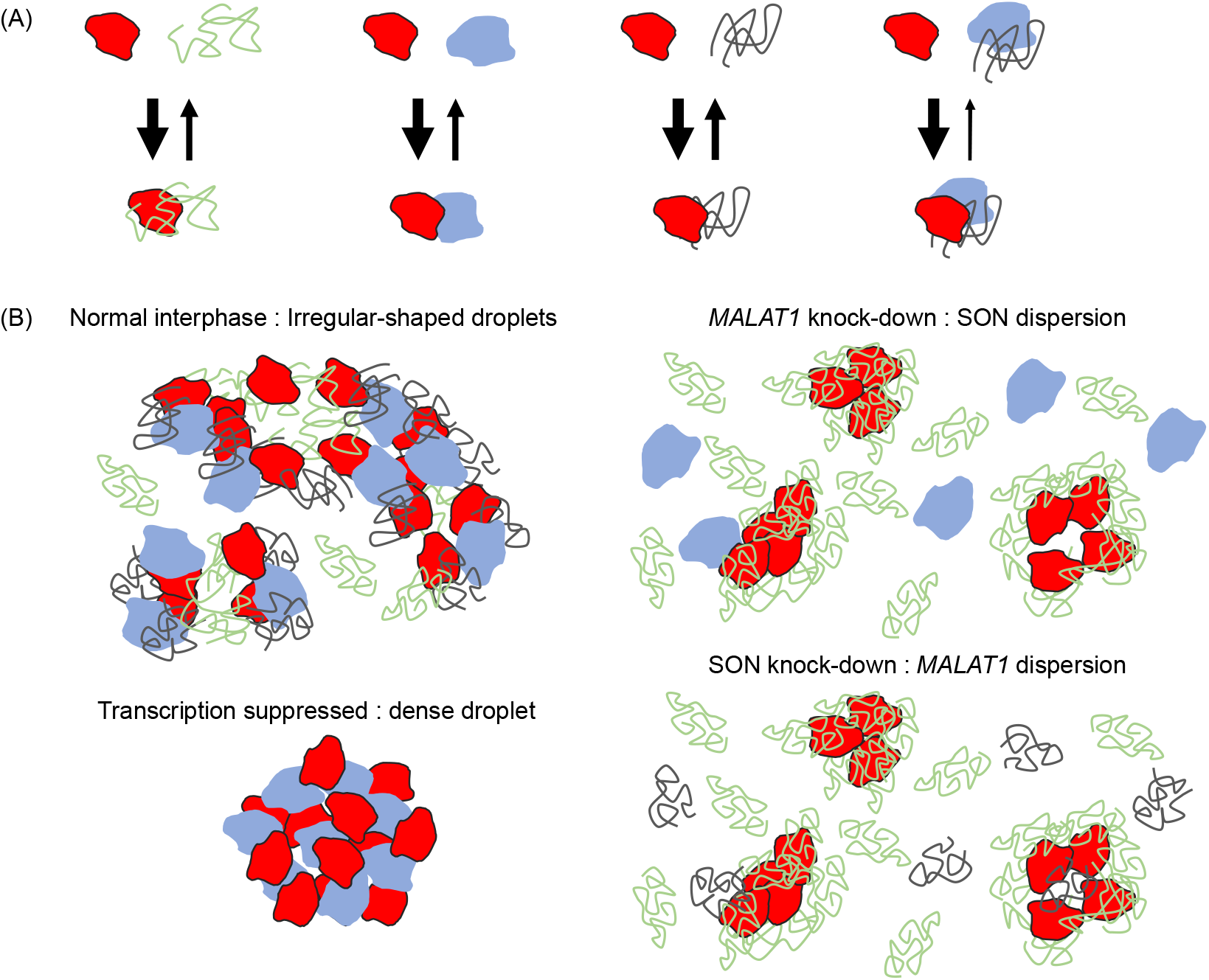
(A) Illustrations of affinities among SON (blue), SRRM2 (red), MALAT1 (black), and pre-mRNA (green). (B) Mechanism and molecular roles of morphological changes in molecular accumulations in local region of nuclear speckles (B). Each molecule color indicates molecular species as explained in Figure 2, and arrow width represents transition rates.

I. SRRM2 exhibited attractive interactions with all other representative molecules of nuclear speckle components; SRRM2-SON interaction was assumed similar (Figure 3–4) or slightly weaker than SRRM2-pre-mRNA interaction (Supplementary Figure S4), SRRM2-MALAT1 interaction was assumed slightly weaker than SRRM2-SON (Figure 7A), and attractive interactions also occurred between SON and *MALAT1*.
II. In the normal interphase of the cell nucleus, pre-mRNAs and SON-*MALAT1* complexes are considered to cluster around SRRM2. However, they were expected to spread spatially owing to their mutual repulsive forces. Notably, the attractive interaction between SRRM2 and SON or *MALAT1* was similar or slightly weaker than that between SRRM2 and pre-mRNA. However, to dissociate from the SON-MALAT1 complex, SRRM2 must dissociate from SON and MALAT1 almost simultaneously, so that the attractive interaction between SRRM2 and SON-MALAT1 complex could be more effective than that between SRRM2 and pre-mRNA (Figure 7A). Therefore, SRRM2 distributed to connect them, which is why the nuclear speckle component was broadly distributed (Figure 7). In such a broad molecular assembly, SON tended to be distributed nearer to *MALAT1*, because the attractive force between SON and SRRM2 was stronger than that between SRRM2 and *MALAT1*.
III. When transcription of the entire nucleus was suppressed, few RNAs existed in the nucleus. Therefore, SON and SRRM2 formed larger dense condensates owing to the mutual attractive interaction between them, without competition between SON and pre-mRNA (Figure 7).
IV. When *MALAT1* was knocked down, SON and pre-mRNA competed for binding with SRRM2. However, more pre-mRNA could bind to SRRM2 than SON even if the affinity of SON to SRRM2 was similar or slightly weaker than that of pre-mRNA. This is because the volume fraction of pre-mRNA was much larger than that of SON; hence, SRRM2 tended to interact with pre-mRNA more frequently than SON. The results showed that most of the SRRM2 was bound to pre-mRNA, while SON was excluded; therefore, SRRM2 sustained its condensate but SON dispersed spatially. Similarly, SRRM2 sustained its condensate, but *MALAT1* dispersed spatially when SON was knocked down (Figure 7).

In the present model, RNAs are assumed to be random polymers. However, RNAs may exhibit various metastable partial folding structures containing many swinging double-strand branches. Additionally, the coupling of some proteins with RNAs may also modify the shape and effective size of RNAs. In these cases, the RNA model can be modified to include slightly smaller and more rigid particles than the presented model. However, the collisional behaviors among such particles are expected to be qualitatively similar to those among large and soft particles in simulations with an appropriate range of *k*_*B*_*T*. Therefore, the results are expected to remain unchanged even after such modifications to the model.

In this study, the effective interactions among the representative components of condensates in nuclear speckles were conjectured to be different from those expected from electrostatic interactions. Thus, these components were expected to form a specific affinity interaction network stronger than electrostatic interactions. However, the chemo-mechanical origins of these interactions cannot be clarified by present arguments. Therefore, further experimental studies are warranted to elucidate the specific molecular binding sites and binding manner of proteins.

Further, the influences of protein modifications including serine phosphorylation of SR proteins and the possibility of interactions should be clarified via other molecules. Moreover, it should be noted that we only simulated a small part of the nuclear speckle with a few molecular condensates. Based on the proposed model with more particles, the simulation of whole nuclear morphological behaviors of nuclear speckles using higher performance computations is a crucial future issue.

## Conclusion

Overall, a coarse-grained molecular dynamics model of nuclear speckle was developed, and the interactions among representative molecules reproduced a rich variety of experimentally observed morphological changes in the droplet-like condensates of this nuclear body. The application of such arguments is expected to provide important insights into the mechanisms underlying the cell state- and cell cycle-dependent behaviors of other nuclear bodies.

## Conflict of Interest

The authors declare that they have no conflict of interests.

## Author Contributions

S.W., N.S., and A.A. conceived and designed the study; S.W. and A.A. conducted the mathematical model construction and simulations; S.W. and A.A. analyzed the data; S.W., N.S., and A.A. wrote the manuscript.

## Data Availability

The evidence data generated and/or analyzed during the current study are available from the corresponding author on reasonable request.

## Acknowledgments

We thank Masashi Fujii for his guidance. This work was supported by Grants-in-Aid for Scientific Research (KAKENHI) [Grant Number 21K06124 (A.A.), 18H05531 (N.S.)] from the Japan Society for Promotion of Science. Computations were partially performed on the NIG supercomputer at ROIS National Institute of Genetics.

---

**Supplementary Figure S1.**
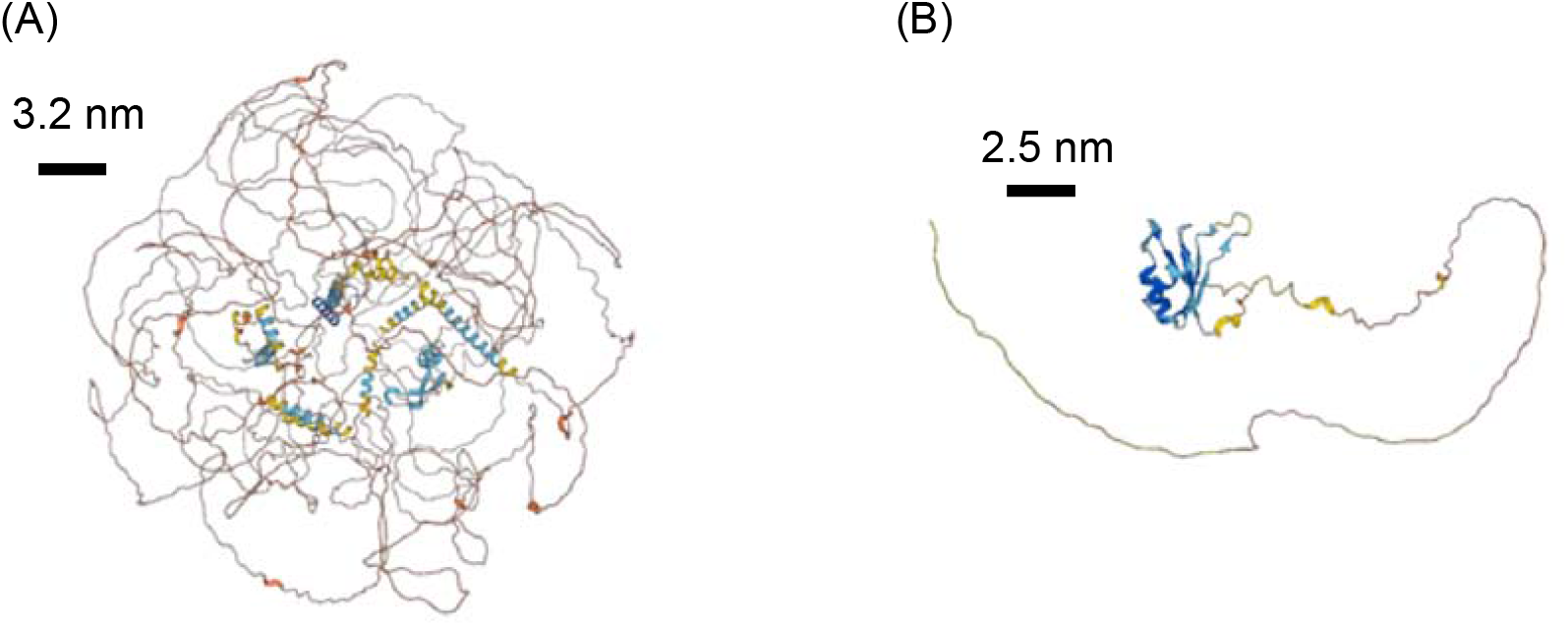
Ribbon descriptions of rough molecular shapes of (A) SON and (B) SRSF2 obtained from UniProt database [27]: ID P18583 and Q01130. The placement of secondary structures on the amino acid sequence is derived from these data. It would be natural to consider that the IDR is distributed more spatially than in these figures.

**Supplementary Figure S2.**
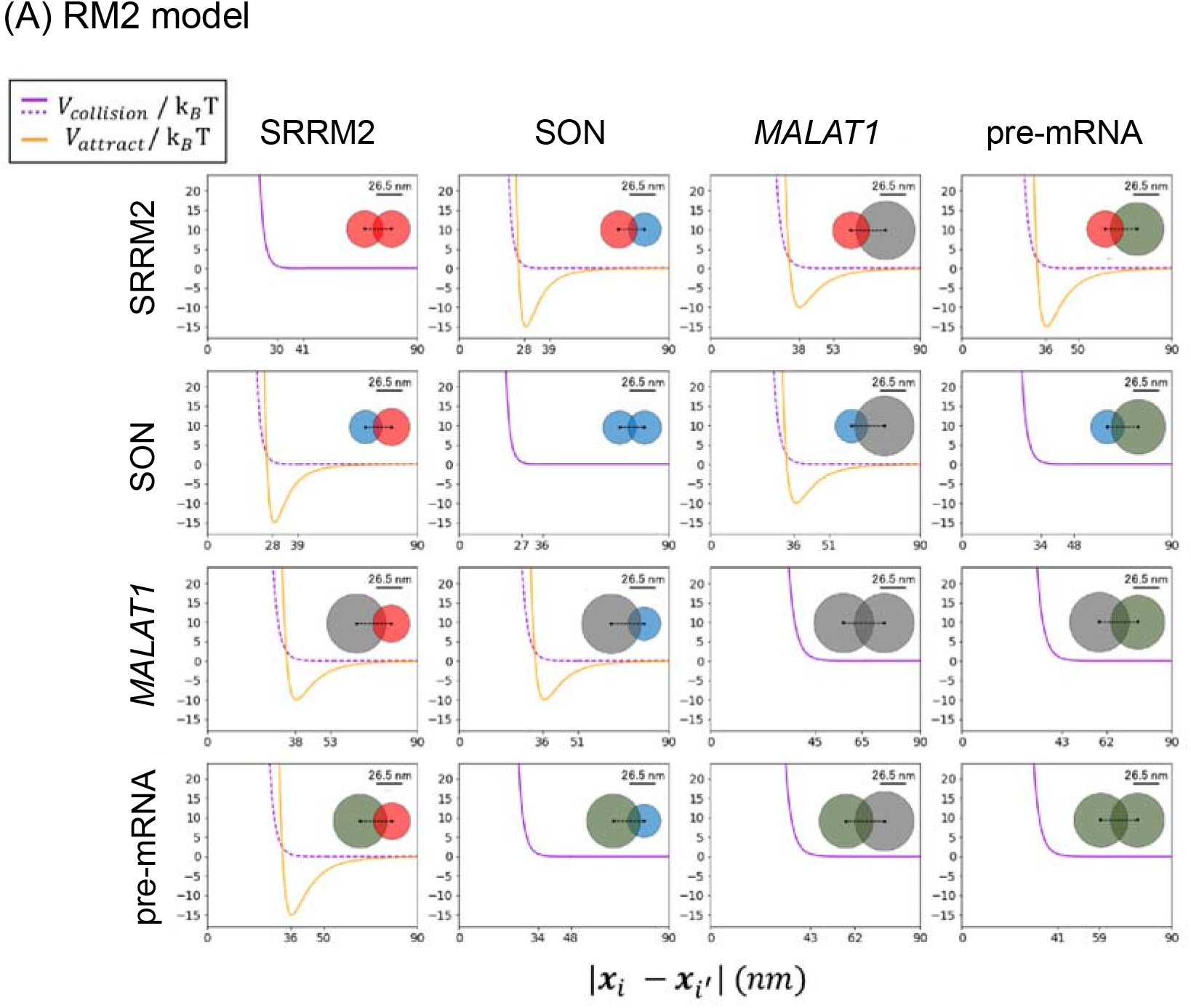
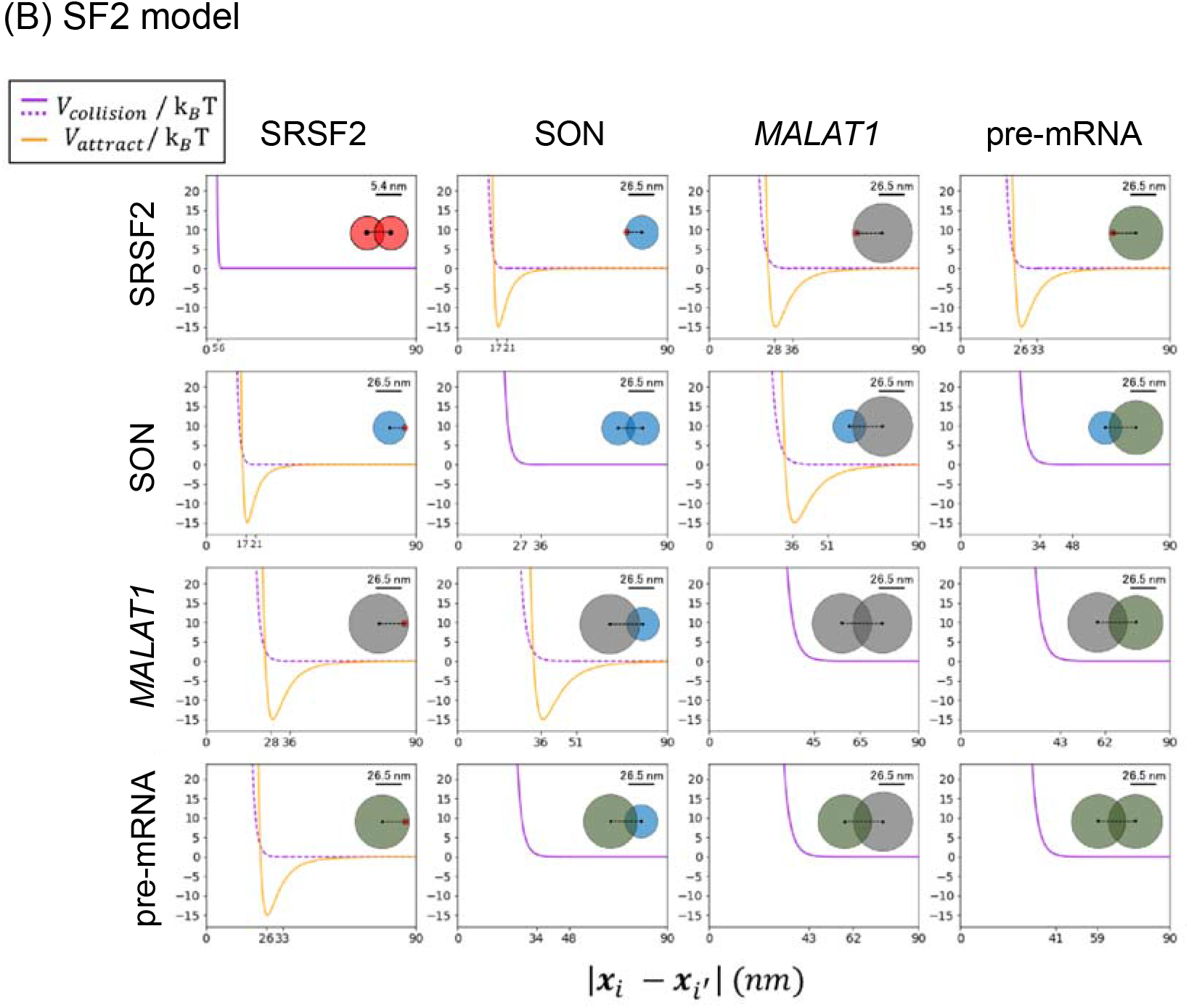
Profiles of interaction potential between each pair of particles in (A) RM2 model and (B) SF2 model that correspond to the interaction network shown in Figure 3. Solid curve indicates each potential profile and broken curve indicates the function 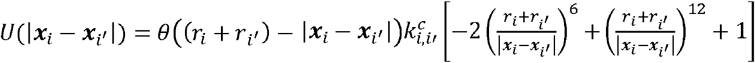. Accordingly, *d*_*i,i′*_ = |***x***_*i*_ − ***x***_*i′*_|, satisfying 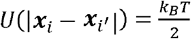 as eqn. (6) in the text. Each pair of two overlapped circles indicates the position of two particles when *d*_*i,i′*_ = |***x***_*i*_ − ***x***_*i′*_|. Color of each particle indicates molecular species as explained in Figure 2.

**Supplementary Figure S3.**
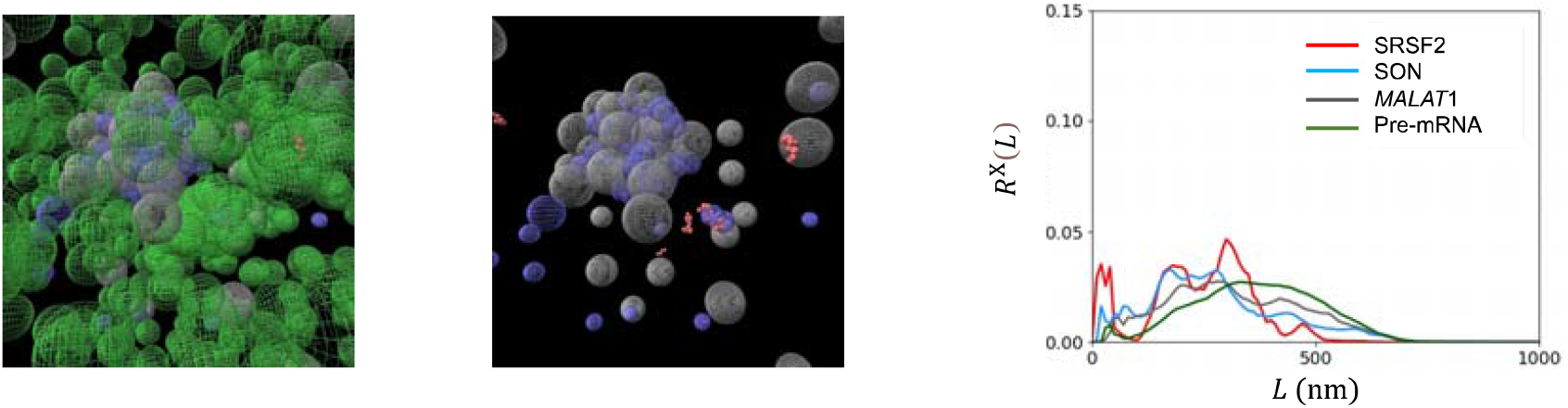
Simulation results of the SF2 model where the attractive strength parameters between SRSF2 and *MALAT1* particles and between SON and *MALAT1* particles was modified uniformly as 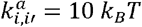. Matrices of interaction networks among particles was given as the same as Figure 3A except that SRRM2 was replaced to SRSF2. Snapshots of the simulations (left), the same snapshots but without pre-mRNA (middle),, and radial distributions of particles, *R*^SRSF2^(*L*), *R*^SON^(*L*), *R*^*MALATI*^(*L*), and *R*^pre−mRNA^(*L*) (right) in the case of normal interphase. Each particle color indicates molecular species as explained in Figure 2. In this case, particles tend to be divided into SRSF2-pre-mRNA particles condensates and SON-*MALAT1* particles condensates with pre-mRNA differently from the results in Figure 4–5.

**Supplementary Figure S4.**
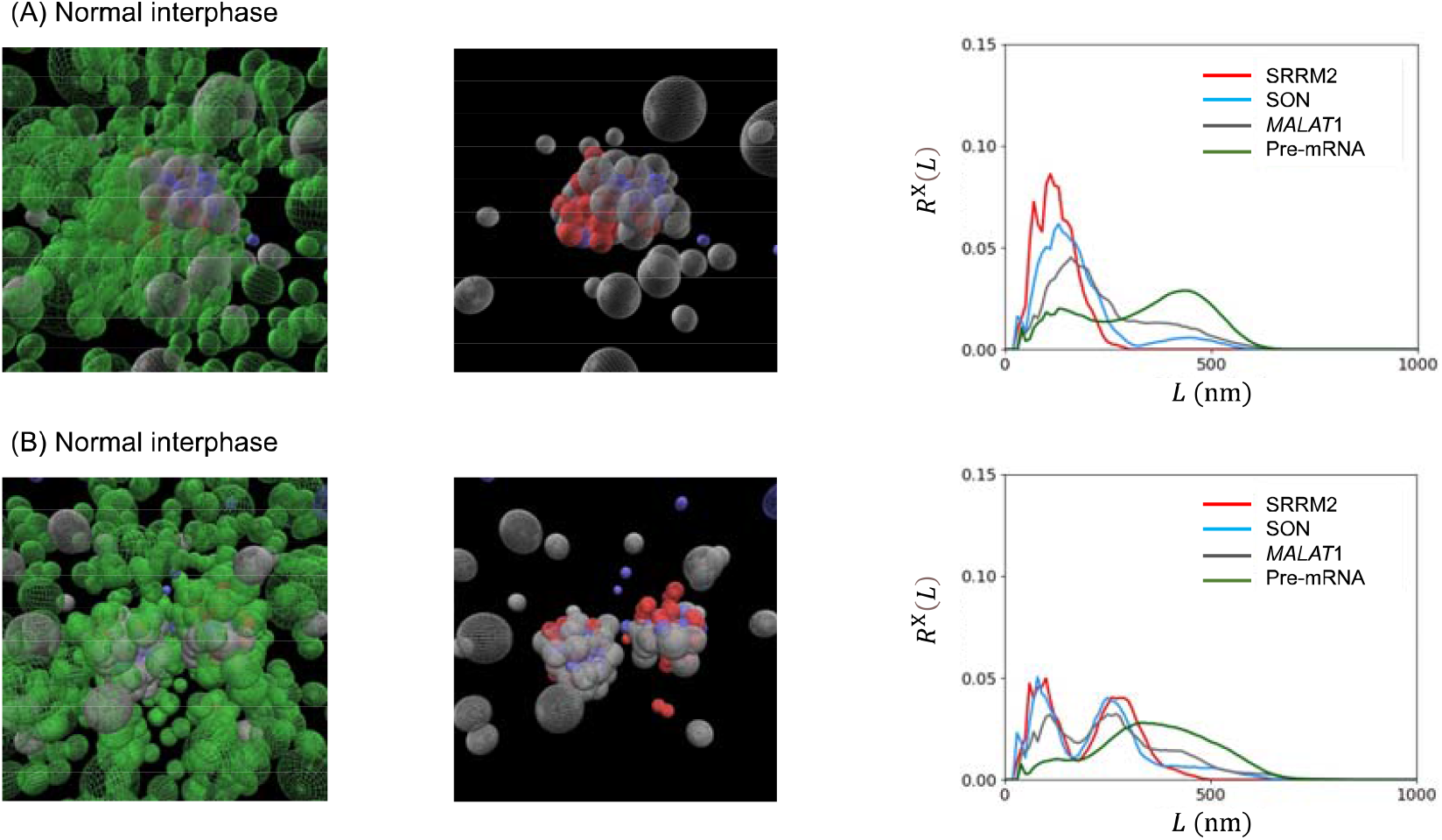
Simulation results of the RM2 model with appropriately designed parameters with a modification that the attractive strength parameter between SRRM2 and pre-mRNA was assumed 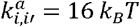. In this case, the attractive force between SRRM2 and SON was slightly weaker than that between SRRM2 and pre-mRNA. (A–B) Simulation results from different initial conditions. Snapshots of the simulations (left) and the same snapshots but without pre-mRNA (middle) in the case of normal interphase. Each particle color indicates molecular species as explained in Figure 2. Qualitatively the same results as in Figure 3 were obtained in vast cases (A), but two particle condensates forming core-shell like structures were also observed in some cases (B). In latter case, the radial distributions of particles, *R*^SRRM2^(*L*), *R*^SON^(*L*), *R*^*MALAT*^(*L*), and *R*^pre−mRNA^(*L*) (right) exhibited two peaks where *R*^*SON*^(*dist*), *R*^*SRRM2*^(*dist*), and *R*^*MALAT*^(*dist*) exhibited peaks at the similar values of *dist*, but *R*^*MALAT*^(*dist*) involved lower peaks and longer tail than *R*^*SON*^(*dist*) and *R*^*SRRM*^ (*dist*).

## Notes

### Competing Interest Statement

The authors have declared no competing interest.

